# Epigenetic liquid biopsies reveal elevated vascular endothelial cell turnover and erythropoiesis in asymptomatic COVID-19 patients

**DOI:** 10.1101/2023.07.28.550957

**Authors:** Roni Ben-Ami, Netanel Loyfer, Eden Cohen, Gavriel Fialkoff, Israa Sharkia, Naama Bogot, Danit Kochan, George Kalak, Amir Jarjoui, Chen Chen-Shuali, Hava Azulai, Hezi Barhoum, Nissim Arish, Moshe M Greenberger, David Vellema, Ramzi Kurd, Eli Ben Chetrit, Davina Bohm, Talya Wolak, Ahmad Quteineh, Gordon Cann, Benjamin Glaser, Nir Friedman, Tommy Kaplan, Ruth Shemer, Ariel Rokach, Yuval Dor

## Abstract

The full spectrum of tissues affected by SARS-CoV-2 infection is crucial for deciphering the heterogenous clinical course of COVID-19. Here, we analyzed DNA methylation and histone modification patterns in circulating chromatin to assess cell type-specific turnover in severe and asymptomatic COVID-19 patients, in relation to clinical outcome. Patients with severe COVID-19 had a massive elevation of circulating cell-free DNA (cfDNA) levels, which originated in lung epithelial cells, cardiomyocytes, vascular endothelial cells and erythroblasts, suggesting increased cell death or turnover in these tissues. The immune response to infection was reflected by elevated B cell and monocyte/macrophage cfDNA levels, and by evidence of an interferon response in cells prior to cfDNA release. Strikingly, monocyte/macrophage cfDNA levels (but not monocyte counts), as well as lung epithelium cfDNA and vascular endothelial cfDNA, predicted clinical deterioration and duration of hospitalization. Asymptomatic patients had elevated levels of immune-derived cfDNA but did not show evidence of pulmonary or cardiac damage. Surprisingly, these patients showed elevated levels of vascular endothelial cell and erythroblast cfDNA, suggesting that sub-clinical vascular and erythrocyte turnover are universal features of COVID-19, independent of disease severity. Epigenetic liquid biopsies provide non-invasive means of monitoring COVID-19 patients, and reveal sub-clinical vascular damage and red blood cell turnover.

## Introduction

SARS-Cov-2 infection directly injures the respiratory system, and inflicts damage to multiple additional tissues including the heart (1), blood vessels (2), pancreas (3), kidneys (4), and liver (5). In most of these cases it is not clear if the damage is caused directly by the viral infection, or whether tissues suffer damage due to the massive host’s immune response. This immune response involves both the innate and adaptive arms of the system (6, 7), and is protective in most cases but can often cause severe inflammatory damage (8). Despite the extensive information that has accumulated since the emergence of the COVID-19 pandemic, the full spectrum of tissues affected by the infection is still not clear. This may partly explain why we still lack tools to predict the clinical course of disease in infected individuals, which ranges from fatal lung failure to a completely asymptomatic presentation. Clinical heterogeneity during the acute phase can potentially account for variation in late-onset clinical phenotypes such as cardiovascular complications (9), new-onset diabetes (10), and the diverse array of long-covid symptoms.

Cell-free chromatin fragments released from dying cells contain extensive information on the identity of cells that released these fragments, and on gene expression programs that operated in the cells prior to their death. In methylation-based liquid biopsies, the tissue origins of cfDNA are inferred based on its tissue-specific methylation patterns. Since the half-life of cfDNA is extremely short (estimated at 15-120 minutes) (11), such analysis can determine the rate of cell death or turnover in specific host tissues close to the time of sampling. Two previous studies characterized the methylome of cfDNA in COVID-19 patients, and deconvoluted it using a partial human cell-type methylome reference atlas (12, 13). The key finding of these studies was that erythroblast turnover is elevated in severe COVID-19 patients and predicts mortality. However, while methylome deconvolution studies can provide an unbiased overview of cfDNA composition, they depend on the quality and breadth of the reference atlas and are typically limited in sensitivity such that tissue contributions to cfDNA amounting to <1% of the total are not detected. To overcome this limitation, we employed deep whole genome bisulfite sequencing (WGBS) of plasma cfDNA followed by deconvolution using a novel extensive human methylome atlas (14). In addition, we used a highly sensitive targeted panel of tissue-specific methylation markers to assess tissue turnover in a cohort of COVID-19 patients including severe hospitalized and asymptomatic patients. Finally, we employed a novel method for cell-free chromatin followed by immunoprecipitation (cfChip-seq) (15) to assess gene expression programs in cells of COVID-19 patients prior to cfDNA release.

## Results

### Patients with severe COVID-19 have a higher concentration of cfDNA, originating in affected tissues

Hospitalized COVID-19 patients (n=120, total 142 plasma samples) had a dramatic, 10- fold elevation in the concentration of cfDNA (**Figure 1A**), suggesting massive cell death. To identify the tissues responsible for the release of cfDNA we employed initially an unbiased approach – deep WGBS (57x coverage in average) on six plasma samples obtained from hospitalized, severe patients infected during summer 2020 and six samples from age-and sex-matched healthy controls. All samples came from unvaccinated individuals.

**Figure 1:**
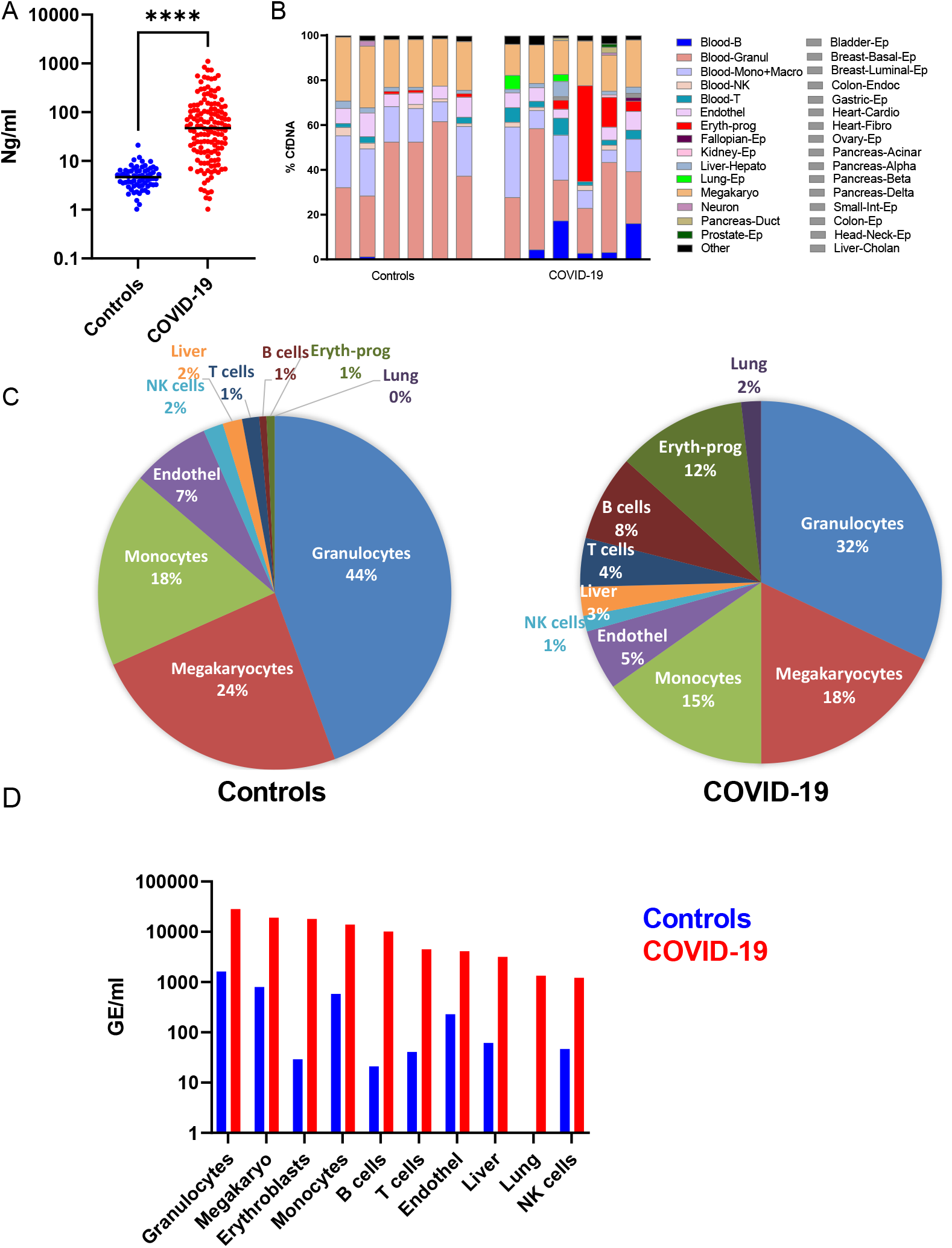
**Whole-Genome Bisulfite Sequencing analysis of severe COVID-19 patients and sex & age matched controls.** (A) CfDNA concentration of hospitalized patients (n=120) and controls (N=68). (B) Deconvolution of plasma samples from severe patients (N=6) and matched controls (N=6). Each column is a single sample. These samples are used for further analysis in the next panels. (C) The average relative contribution of indicated cell types in patients and controls. (D) Absolute concentration of cfDNA from indicated cell types in patients and controls, derived by multiplying the fraction of cell type-specific cfDNA by the total concentration of cfDNA in the sample. Values are expressed as genome equivalents per ml plasma. \**P* < 0.05, \*\**P* < 0.01, and \*\*\**P* < 0.001 ****P<0.0001. Band indicates the median value.

Deep WGBS was followed by unsupervised deconvolution based on an extensive reference atlas of human tissue-specific methylomes of 40 cell types (14), allowing highly accurate identification of methylation patterns derived from all these potential sources (14). The composition of cfDNA in the healthy controls is consistent with previous studies. Blood cell types account for 91% of the cfDNA (granulocytes 44%, megakaryocytes 24%, monocytes/macrophages 18%, NK cells 2%, and B cells, T cells and erythroblasts each contributing ∼1%), and the rest originated from vascular endothelial cells (7%) and hepatocytes (2%) (**Figure 1B,C**). Consistent with previous reports (16, 17) we do not find evidence of DNA derived from the lung in healthy individuals. In contrast, COVID-19 patients had an elevation in the fraction of cfDNA that originated from the liver (3%), T cells (4%), B cells (8%), erythroblasts (12%) and lungs (2%), at the expense of the relative contribution of granulocytes (mainly neutrophils), megakaryocytes and monocytes/macrophages (32%, 18% and 15%, respectively) (**Figure 1B,C)**. When considering both the relative contribution of each tissue and the total concentration of cfDNA, we observed an elevation in the concentration of cfDNA derived from all blood lineage, in addition to vascular endothelial cells, hepatocytes and lung epithelial cells (**Figure 1D**).

These findings provide an unbiased view of cell turnover in severe COVID-19, which is consistent with previous reports and with the clinical nature of the disease. Elevated lung cfDNA reflects excessive lung cell death, which is expected given that the lung is the primary site of infection and the key contributor to clinical damage. Erythroblast cfDNA was previously shown to be elevated in COVID-19 patients, although it was suggested that only morbidly sick patients show this phenomenon, while we identify a massive elevation in erythrocyte cfDNA in 4/6 patients (see below further details using targeted analysis). Similarly, elevated cfDNA from immune cells is to be expected, as a reflection of the normal immune response of the host. Vascular endothelium-derived cfDNA may reflect the documented evidence of vascular damage in patients with severe COVID-19 (18).

### A targeted assay for cfDNA biomarkers relevant to COVID-19 patients

To further characterize cfDNA content in COVID-19 patients, we applied a targeted cfDNA methylation assay on the entire cohort of plasma samples. This allows for ultra-deep assessment of specific targets at a modest cost. The use of targeted markers allows to sequence essentially every molecule from a given marker locus, providing higher sensitivity without compromising specificity. We designed a panel of 37 methylation markers, containing loci that are specifically unmethylated or specifically methylated in cell types relevant to COVID-19 patients. The panel contained markers of specific immune cell types: B-cells (4 independent markers), T-cells (2), CD8+ T-cells (1), NK cells (2) monocytes/macrophages (3) and neutrophils (3). In addition, it included methylation markers of other relevant blood cell types including erythroblasts (5) and megakaryocytes (2), as well as markers of lung epithelial cells (6), vascular endothelial cells (4) and cardiomyocytes (5). We validated the specificity of these markers (**Supplementary Figure S1**) and established a protocol for two multiplex PCR conditions that could amplify all markers from the cfDNA extracted from each plasma sample (see Methods).

With this targeted assay at hand, we moved to assessment of samples from the larger cohort of 120 hospitalized patients and 40 controls, all of whom were unvaccinated. Consistent with the methylome deconvolution analysis, we observed a highly significant increase in cfDNA originating in immune cells, including elevated levels of cfDNA from B cells, T cells, monocyte/macrophages, neutrophils, CD8 T cells and NK cells (**Figure 2A**). In addition, cfDNA of erythroblasts and megakaryocytes was dramatically elevated, reflecting increased turnover and production of red blood cells and platelets (**Figure 2B**).

**Figure 2:**
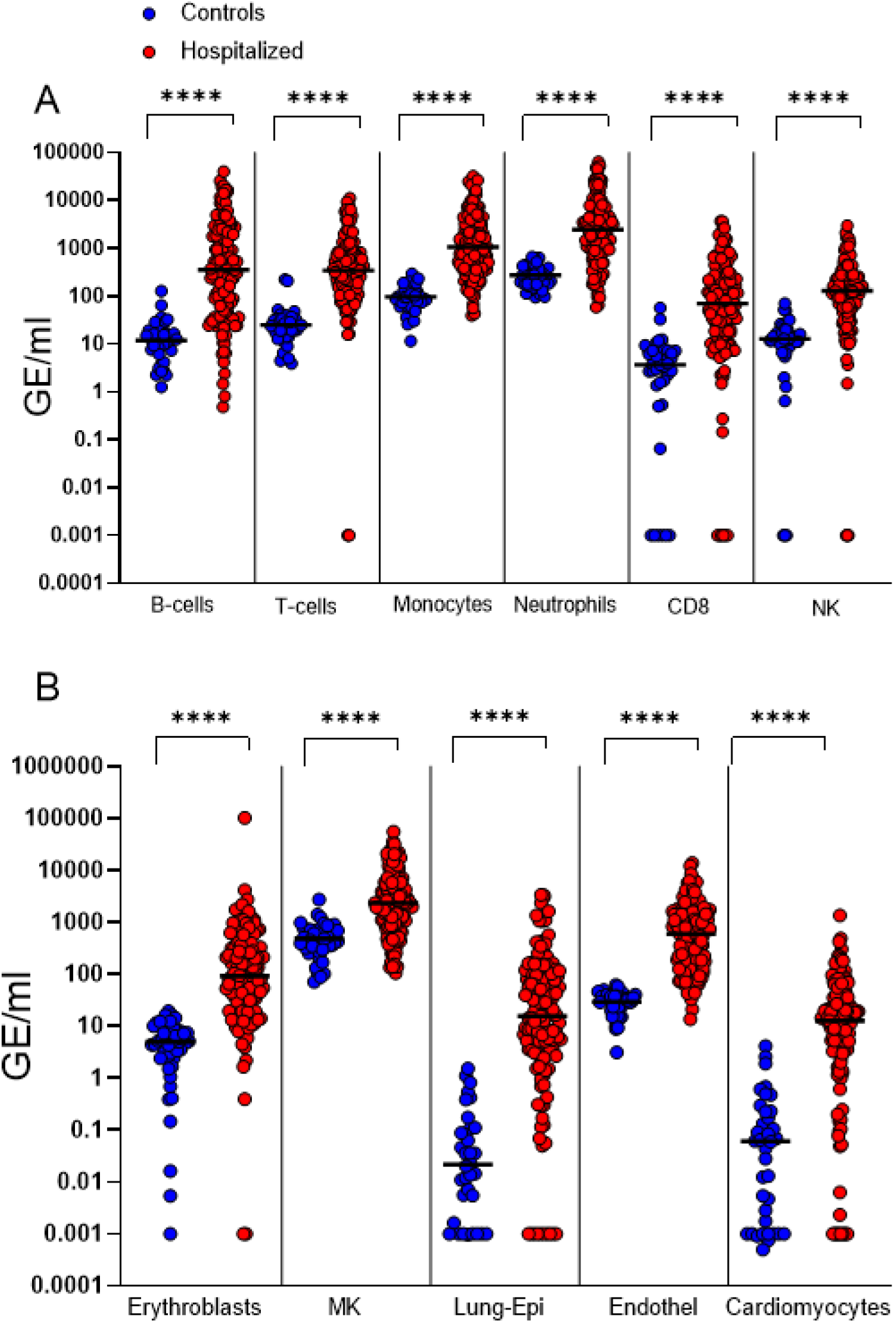
Targeted analyses of cfDNA origins in hospitalized patients. Each dot represents a sample. In total, the analysis includes 120 hospitalized patients and 30-45 healthy controls. **(A)** Immune-derived cfDNA markers. **(B)** cfDNA methylation markers of erythroblasts, megakaryocytes (MK), lung epithelial cells, vascular endothelial cells and cardiomyocytes. Note that the median level of cardiomyocyte cfDNA is 12 GE/ml, potentially explaining why this was not detected by the less-sensitive deconvolution analysis. \**P* < 0.05, \*\**P* < 0.01, and \*\*\**P* < 0.001 ****P<0.0001. Horizontal lines indicate the median value.

We also noticed elevated levels of cfDNA derived from lung epithelial cells and vascular endothelial cells, consistent with the deconvolution analysis (**Figure 2B**). Importantly, some (but not all) patients had significantly elevated levels of cardiomyocyte cfDNA, suggesting ongoing cardiac damage (**Figure 2B**), consistent with reports on heart damage in severe COVID-19 patients (1, 19).

Thus, targeted analysis of cfDNA in hospitalized COVID-19 patients reveals an extensive immune response, and elevated turnover of red blood cells, megakaryocytes, lung epithelial cells and vascular endothelial cells. In addition, the high sensitivity of the targeted assay showed striking evidence of cardiomyocyte death in hospitalized patients.

### cfDNA correlates to clinical severity score

To understand the potential of cell type-specific cfDNA to explain clinical phenomena, we generated a correlation matrix quantifying the relationship between cfDNA parameters and key clinical records for the entire set of 120 patients hospitalized with COVID-19. The matrix included age and sex, standard biochemical markers, blood cell counts and the WHO progression scale of COVID-19 clinical severity (20), as well as cfDNA parameters.

To verify internal consistency, we first determined the correlation between non-cfDNA parameters. Consistent with previous studies (21, 22), clinical severity was correlated with neutrophil counts, and negatively correlated with lymphocyte counts (i.e. associated with lymphopenia). Biochemical measures of stress and inflammation (CRP, D-Dimer and ferritin) also correlated with clinical severity of COVID-19, as well as high neutrophil counts and low lymphocyte counts (**Figure 3**).

**Figure 3:**
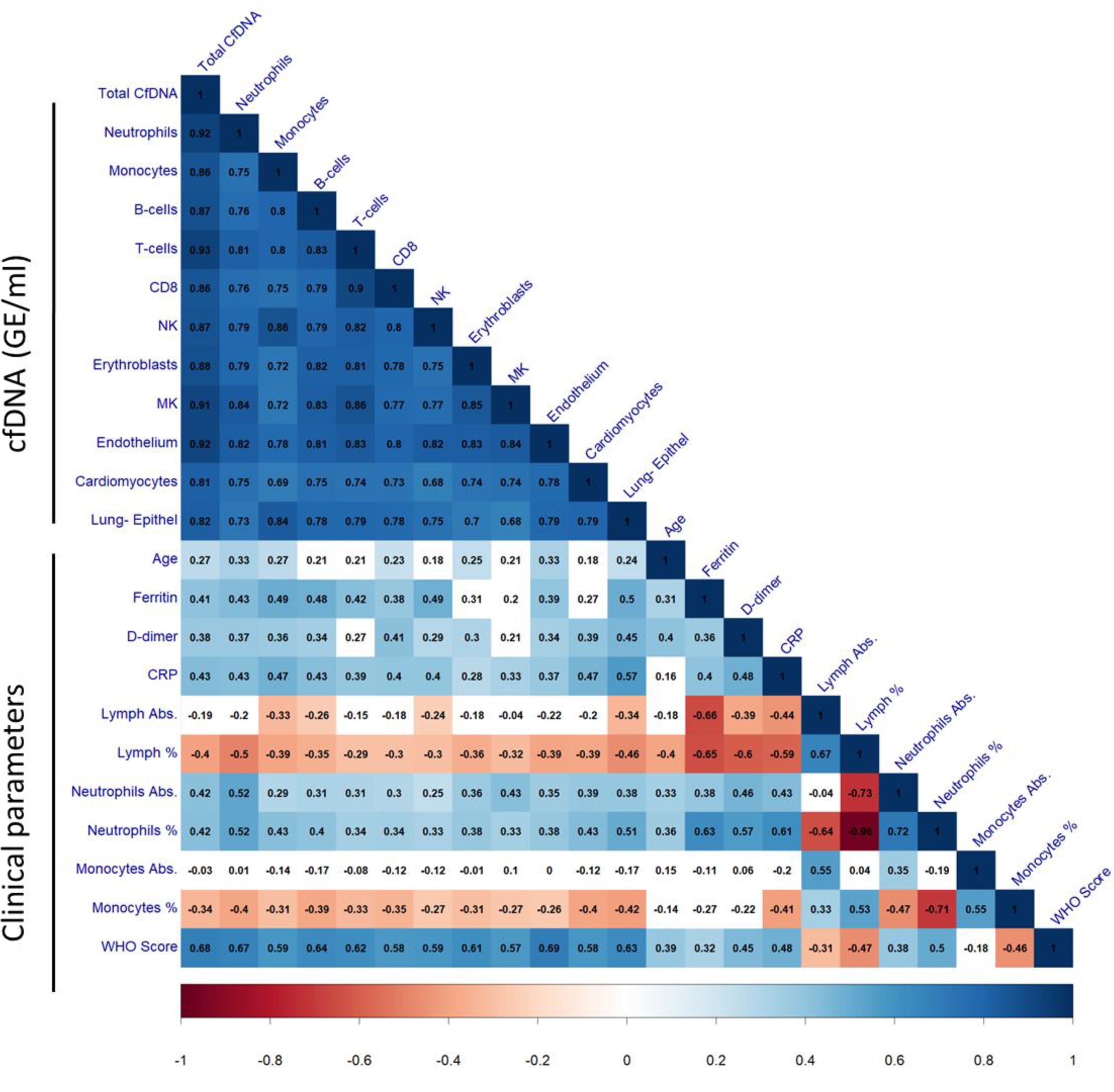
Correlations between cell type-specific cfDNA, clinical and laboratory parameters, and WHO clinical score. Correlation coefficients are presented in black inside each square. Only statistically significant correlations (q<0.05) are colored, and the blue-white-red color scale reflects of coefficient value.

The concentration of cfDNA from immune cells, erythroblasts, heart, lung and vascular endothelial cells was positively and significantly correlated to disease severity, measured by the WHO score at the time of sampling. Vascular endothelial cell cfDNA showed the strongest correlation to disease severity, followed by the total concentration of cfDNA, neutrophil cfDNA, B cell cfDNA and lung epithelium cfDNA (**Figure 3**). Note that the correlation of cfDNA to disease severity was generally much stronger than the correlation of circulating blood cell counts and all biochemical markers to disease severity, highlighting the direct relationship between clinical condition and tissue damage as reflected by cfDNA.

### CfDNA as a prognostic marker for clinical course

To assess the utility of tissue-specific cfDNA in predicting the clinical trajectory of COVID- 19, we generated a simple score of future clinical course. This score considers the WHO score at the day of plasma sampling, and the maximal score the patient reached afterwards during hospitalization (until discharge or death). With this, each individual sample is assigned a “recovering” or “deteriorating” status based on the clinical course of the patient.

As shown in **Figure 4A**, hospitalized patients that have clinically deteriorated in the days after blood sampling had at the time of sampling significantly higher levels of cfDNA from multiple immune cell types including neutrophils, monocyte/macrophages and NK cells, and to a lesser extent T cells. Interestingly, B cell cfDNA was not a predictor of clinical outcome, despite B cells being a critical determinant of humoral immunity. This finding, along with the weak correlation between T cell cfDNA and clinical outcome, suggests that deterioration of hospitalized patients is dictated mostly by performance of their innate immune system, combined with the extent of damage to vital organs (see below) (6, 23).

**Figure 4:**
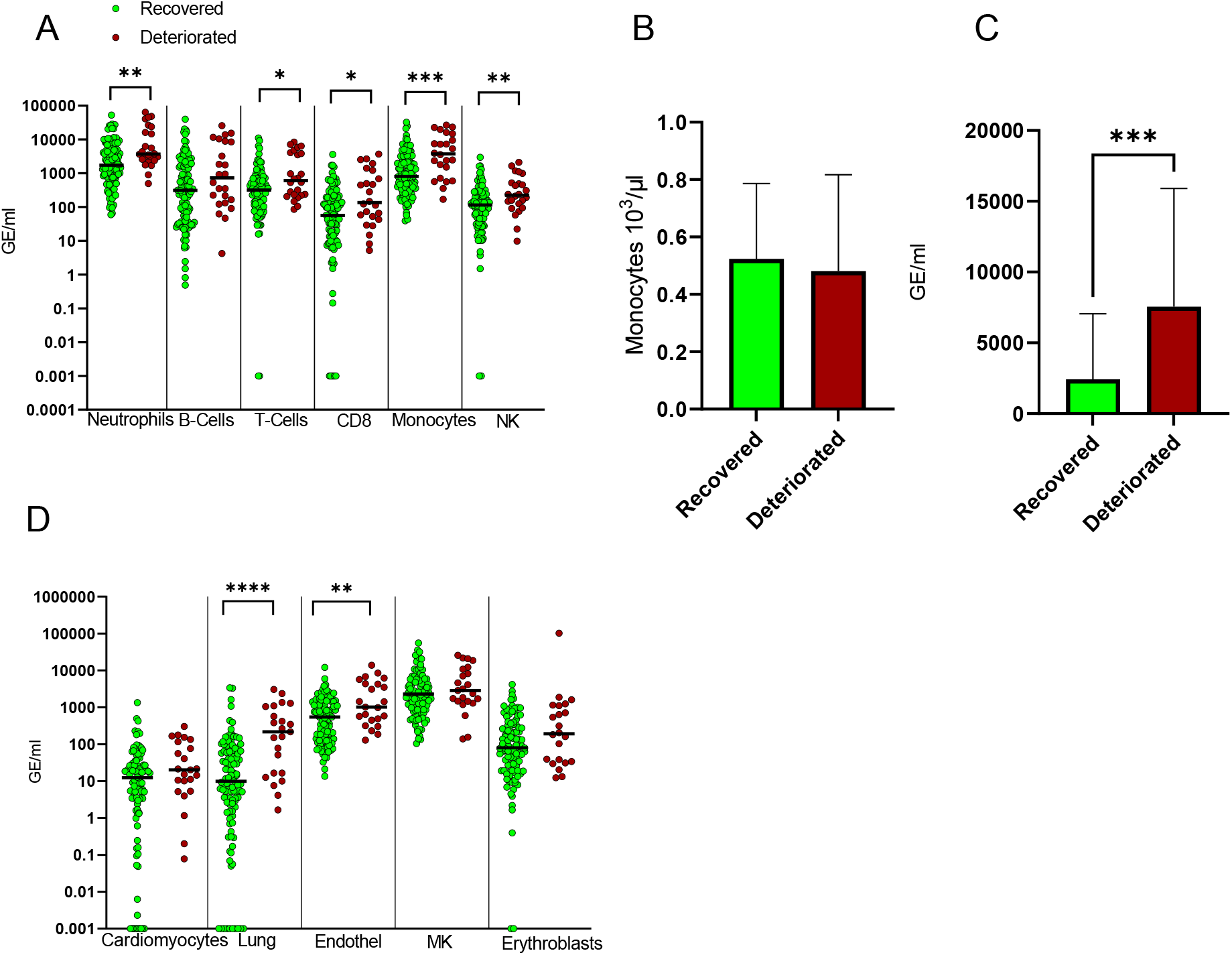
**Correlation between cell type-specific cfDNA and future clinical course.** WHO score on the day of sampling was subtracted from the maximal WHO score during the hospitalization, following the sampling day. Non-positive values were scored as clinical recovery, while positive values were scored as deterioration. (A) Immune-derived cfDNA levels in patients that recovered or deteriorated clinically after sampling. (B) Monocyte cell counts in patients that deteriorated or recovered. (C) Monocytes/macrophages-derived cfDNA in patients that deteriorated or recovered. (D) cfDNA from cardiomyocytes, lung epithelial cells, vascular endothelial cells, megakaryocytes (MK) and erythroblasts in patients that recovered or deteriorated clinically after sampling. \**P* < 0.05, \*\**P* < 0.01, and \*\*\**P* < 0.001 ****P<0.0001. Band indicates the median value.

The correlation of monocyte cfDNA levels to clinical deterioration was particularly strong. Strikingly, the counts of circulating monocytes were not associated with clinical course, suggesting that monocyte/macrophage cfDNA levels reflect the amount monocyte damage or turnover taking place within tissues (potentially a result of infection, see Discussion), without altering systemic cell counts (**Figure 4B,C**).

Among non-immune cfDNA, lung epithelial cells and vascular endothelial cells were significantly associated with future deterioration of hospitalized patients, while cardiomyocyte, megakaryocyte and erythroblast cfDNA were not. While the correlation between clinical course and the extent of lung damage as reflected in lung cfDNA is expected, the relationship between vascular endothelial cell turnover and clinical outcome is less trivial, and indicates a prominent involvement of vascular damage in the pathogenesis and course of COVID-19 (see below).

### Asymptomatic patients show sub-clinical elevation of vascular endothelial cell turnover and in erythropoiesis

We characterized cfDNA in patients with asymptomatic or mild disease, who donated blood while being quarantined due to a positive SARS-Cov-2 PCR test. This is an under-studied population, which can presumably provide insights into the cfDNA manifestation of a successful immune response, and to uncover sub-clinical damage to organs. The cfDNA concentration in these individuals (n=19 patients, 8.6 ng/ml) was significantly higher than in healthy controls (n=68 donors, 4.7 ng/ml) (p value=0.0001), yet it was far lower than the concentration of cfDNA in the hospitalized patients described above (n=120 patients, n=142 samples, 47.1 ng/ml) (p value=0.0001) (**Figure 5A**).

**Figure 5:**
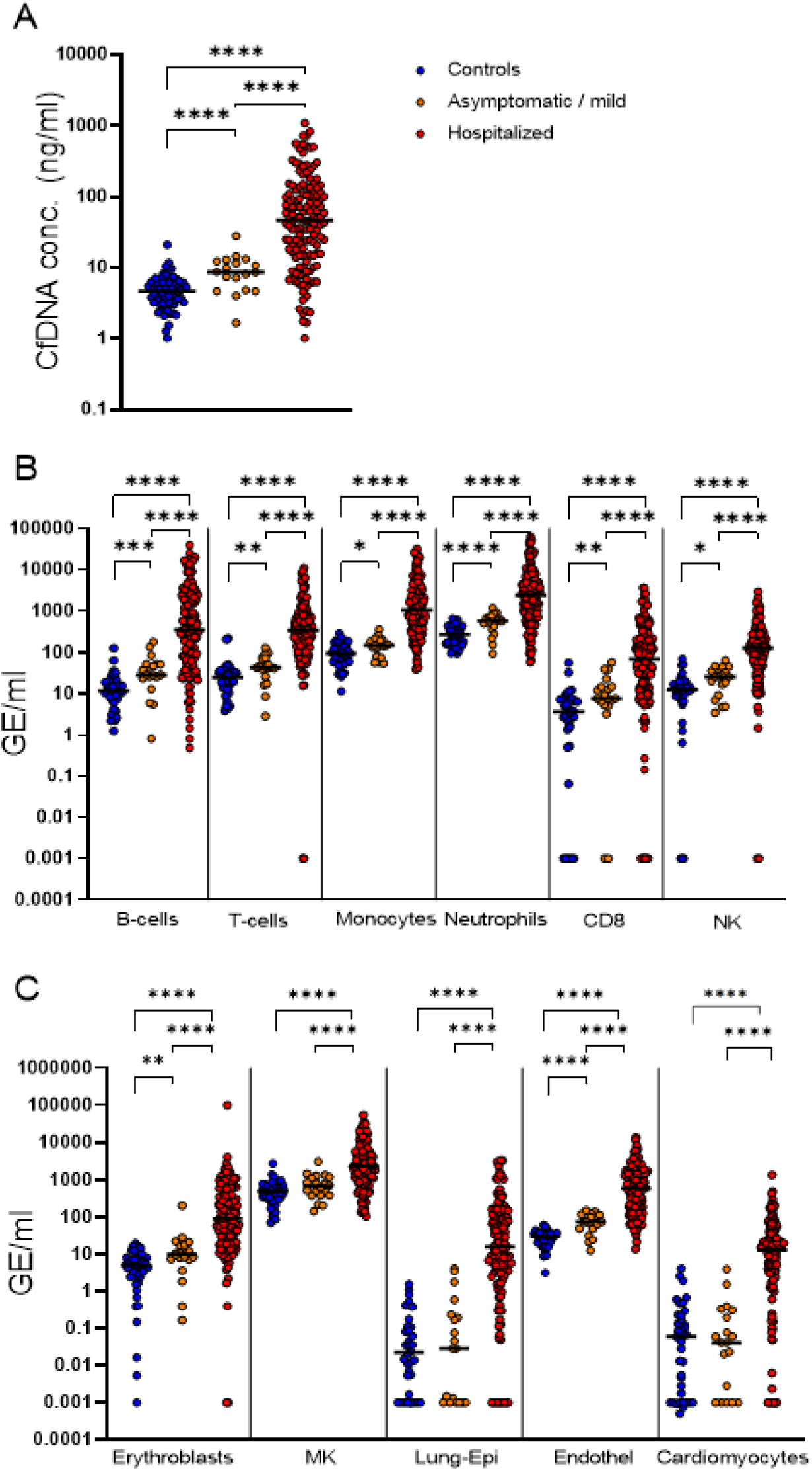
**Cell type-specific cfDNA in patients with asymptomatic or mild COVID-19 (n=19) compared with hospitalized patients (n=120) and healthy controls (n=30-45).** (A) Total cfDNA concentration. (B) Immune-derived cfDNA. (C) cfDNA from erythroblasts, megakaryocytes, lung epithelial cells, vascular endothelial cells and cardiomyocytes. \**P* < 0.05, \*\**P* < 0.01, and \*\*\**P* < 0.001 ****P<0.0001. Band indicates the median value.

**Figure 6:**
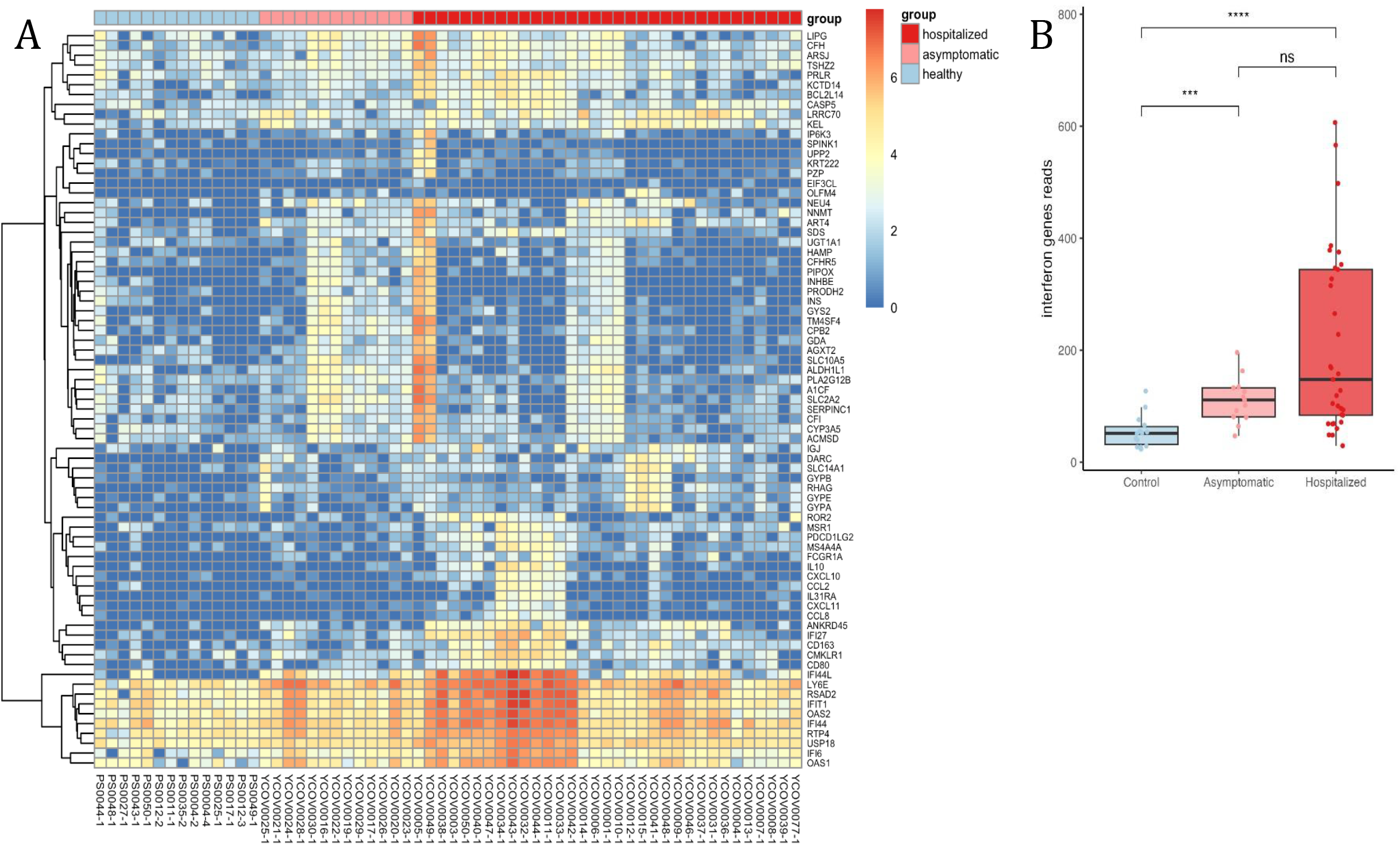
**Chromatin immunoprecipitation of cell-free nucleosomes analysis of COVID-19 patients, asymptomatic and controls.** (A) Unsupervised clustered heat map of the genes that are significantly elevated in the hospitalized COVID-19 patients, compared to controls. (B) reads of genes from a predefined interferon response gene set in the three groups.

The asymptomatic patients had elevated levels of cfDNA from all immune cell types measured, likely reflecting cellular turnover associated with the successful mounting of an immune response to curb the virus (**Figure 5B**). The much higher concentration of immune-derived cfDNA in hospitalized patients reflects either increased activity needed to fight off a higher viral load, or alternatively a pathologic overactivity (24).

Asymptomatic patients did not have higher than normal levels of megakaryocyte cfDNA, lung cfDNA or cardiomyocyte cfDNA. However, these patients showed significantly elevated cfDNA levels derived from erythroblasts and from vascular endothelial cells, even though there was no clinical indication of disruption of red blood cell homeostasis or damage to the vasculature (**Figure 5C**). This finding, in addition to elevation of the d-dimer protein (**Supplementary Figure S5** and previous studies (25, 26)) and previously reported increased counts of nucleated RBC (27), suggests sub-clinical vascular damage compensated by endothelial cell regeneration, as well as enhanced rate of erythropoiesis (see discussion).

### Circulating chromatin indicates interferon response in dying cells

The course of SARS-Cov-2 infection involves major physiological processes including a strong immune response and cytokine storm as mentioned previously. These processes entail the activation of gene expression programs that are inactive in the normal state of cells. We hypothesized that some of these programs occur in the dying cells and might be reflected in cfDNA. In order to test this, we made use of cfChIP-seq, a recently introduced method of chromatin immunoprecipitation of cell-free nucleosomes carrying active chromatin modifications, which allows to infer gene expression in cells undergoing apoptosis, prior to the release of cfDNA (15).

Specifically, we used cfChIP-seq with an antibody against Histone 3 lysine 4 trimethylation (H3K4me3), a chromatin modification that marks open promoters (28, 29). We applied cfChIP-seq to plasma samples from hospitalized COVID-19 patients (n=33), asymptomatic COVID-19 patients (n=13) and a healthy control group (n=14), with an average yield of ∼3.5 million unique reads per sample, which are a genome-wide representation of open promoters, similar to RNA-seq gene expression counts.

Using statistical tests designed for cfChIP-seq we searched for genes with differential coverage between the hospitalized and healthy samples (see Methods). Several dozen genes came up as significantly elevated in the hospitalized group (**Supplemental Figure 2** and **Supplemental table S6**). Inspection of these genes revealed three clusters with distinct behaviors (**Figure 5A**). The first cluster contains genes that are expressed only in the liver (e.g. *SERPINC1*, *CFHR5*) and the entire set of genes is strongly enriched for the liver (see enrichment of gene sets for all clusters in **Supplemental table S7,8,9**). The genes in this cluster were especially high in two of the hospitalized patients but were observed also in other hospitalized and asymptomatic patients, suggesting elevated death of hepatocytes in these cases. The second cluster includes the erythroblast specific glycophorin genes (*GYPA, GYPB, GYPE*), and was observed in a different subset of the hospitalized samples, as well as one of the asymptomatic samples. This signal presumably reflects elevated turnover of red blood cells and increased rate of erythropoiesis, and is consistent with the methylation-based observation of elevated erythroblast cfDNA (**Figure 1,2**).

The third cluster, in contrast to the first two, does not reflect death of a specific cell-type but rather a gene expression program that is typically inactive in cells. The genes in this cluster are enriched for the interferon response and include genes such as *IFIT1*, *IFI6, and IFI27*.

Computing the cumulative signal over a predefined interferon response gene set from the Human MSigDB Collections (30, **Supplemental table S10**), we observed a significant signal in the hospitalized and asymptomatic groups compared to the healthy samples (Wilcox test *P = 8.2X10^-6^* and *P* = 0.00042 respectively), which is particularly pronounced in a subset of samples from hospitalized patients (**Figure 5B**).

Overall, these results uncover an interferon response occurring in the dying cells of COVID-19 patients.

## Discussion

### Unbiased and targeted epigenetic liquid biopsies in COVID-19

Analysis of cell-free DNA methylation and histone modifications allows the characterization of cellular turnover or death taking place inside organs, as well gene expression program operating in cells prior to death and release of cfDNA. In the case of solid organs, this is equivalent to a standard biopsy, with the advantage that information is summed across the entire organ. In the case of the immune cells, cfDNA informs on immune and inflammatory processes taking place within organs, which are not reflected in blood cell counts (31). Here we applied this technology in three forms: an unbiased deconvolution of the plasma methylome, targeted analysis of cell type-specific methylation markers, and cfChip-seq. An unbiased approach to cfDNA methylation of COVID-19 patients was previously applied by two studies (12, 13). Our work differs in several aspects. First, we sequenced plasma at a coverage more than an order of magnitude deeper (57x vs 1.3X (12)). Second, we used a new, more extensive human cell type methylome atlas that allows for more accurate interpretation of cfDNA methylation patterns (14). Third, the targeted assay, while limited to pre-defined loci, allows for a more accurate and sensitive detection of cfDNA from a given source. Fourth, we applied the cfChip-seq assay which provide for the first time information on gene expression programs in COVID-19 patients, specifically in cells that are turning over and releasing cfDNA.

### Tissue dynamics in COVID-19

The analysis of cfDNA methylation and chromatin patterns led to several insights regarding disease dynamics in COVID-19 patients:

i. Evidence of frequent cardiomyocyte death in hospitalized patients. While cardiac damage is a known component of COVID-19 pathology (32), it has not been appreciated that cardiac cell death is a feature shared by most hospitalized patients. We note however that the magnitude of the phenomenon is small (median of 12 cardiac genome equivalents per milliliter plasma is patients). Further study is required to determine the long-term clinical significance of elevated cardiomyocyte cfDNA in hospitalized COVID- 19 patients.
ii. A strong correlation between disease severity and general cfDNA concentration as well as the contribution of specific immune cell types and solid organs. Almost all cfDNA parameters measured showed a statistically significant, positive correlation to the standard WHO score of COVID-19 severity. Notably, the correlation of severity to cfDNA parameters was stronger than the correlation to all standard biomarkers including CRP, ferritin, d-dimer and lymphocyte counts. This underscores the direct relationship between tissue damage and cfDNA. We propose that the short half-life of cfDNA (15-120 minutes, compared with hours or days for standard biomarkers), a feature which makes it an acute biomarker for tissue turnover at the time of sampling, further strengthens the correlation to the current clinical condition of a patient.
iii. Prediction of clinical deterioration by the levels of cfDNA derived from innate immune cells, lung epithelium and vascular endothelial cells. This finding is consistent with extensive literature on the fundamental role of the innate immune system in setting the clinical course of COVID-19. The correlation of monocyte/macrophage cfDNA, but not monocyte counts, to clinical deterioration is particularly intriguing. It is in agreement with a recent report on the prominent role of monocytes in critically ill patients, including the effect of monocyte-expressed genes on vascular permeability in the lung (33). One potential interpretation is that elevated monocyte cfDNA reflects direct viral infection of monocytes (34), which is indicative of viral spread and immune failure. Alternatively, the abnormally high turnover of monocytes may reflect pathogenic activation of the immune system that inflicts tissue damage beyond direct damage due to infection. We acknowledge a current limitation in the specificity of our markers, which are unable to distinguish between DNA originating in circulating monocytes and DNA originating in tissue-resident macrophages. Further refinement of the methylome atlas may resolve this ambiguity.
iv. Heightened interferon response. SARS-Cov-2 infection causes a massive elevation in the expression of interferon-stimulated genes, as measured in leukocytes and in material from invasive biopsies (35–37). The identification of an interferon response in cell-free chromatin underscores the potential of epigenetic liquid biopsies to reveal gene expression program in cells prior to their death. Further refinement of the cfChip-Seq technology (e.g. analysis of chromatin marks characteristic of tissue-specific enhancers) may allow identification of the cell types that have activated the interferon response prior to cell death.
v. Sub-clinical elevation of erythroblast and vascular endothelial cell cfDNA in patients with asymptomatic or mild disease. The elevated levels of erythroblast cfDNA observed in our experiments are consistent with a previous report that erythroblast cfDNA is elevated in pre-morbid COVID-19 patients (12), and support the idea that SARS-Cov-2 infection causes a disruption in erythropoiesis (27). Importantly, the higher sensitivity of targeted assay allowed the detection of mild though significant elevation of erythroblast cfDNA in the majority of patients, including severe patients that eventually survived and, most strikingly, asymptomatic patients. This suggests that SARS-Cov-2 infection causes a profound disruption of erythropoiesis, regardless of disease severity or clinical manifestation. A higher viral load and clinical deterioration may increase further the magnitude of this phenomenon. Since red blood cell counts do not change in most COVID-19 patients (21), elevated levels of erythroblast cfDNA likely mean a higher rate of turnover. We speculate that SARS-Cov-2 infection shortens the lifespan of red blood cells, triggering a compensatory increase in erythropoiesis which is manifested in elevated erythroblast cfDNA. In severe COVID-19, elevated counts of circulating nucleated red blood cells have been reported, potentially accounting for some of the observed elevation in erythroblast cfDNA (27). Additional studies are needed to test this hypothesis and understand the underlying mechanisms. In addition to erythroblast cfDNA, vascular endothelial cell cfDNA levels were significantly elevated in the asymptomatic patients. While vascular damage was shown to take place in a Rhesus Macaque model of COVID-19 (38), evidence in human patients is more limited due to the paucity of circulating biomarkers of vascular endothelial damage. The finding that even patients with asymptomatic disease have elevated levels of vascular endothelial cfDNA suggests that COVID-19 causes a sub-clinical increase of vascular turnover, via either direct viral damage or via immune mediators. As long-covid can develop even in patients that had a relatively mild disease, it is particularly interesting to determine if cfDNA abnormalities in mild cases are prognostic of long-covid.

In summary, epigenetic liquid biopsies open a non-invasive window into tissue dynamics in patients with COVID-19, including sub-clinical damage and dynamic processes that cannot be assessed via existing biomarkers. Further studies of patients with long-covid will reveal the prognostic and diagnostic utility of cell-free DNA and chromatin in this disease. Beyond COVID-19, our findings underscore the potential of epigenetic liquid biopsies to characterize complex pathologies that do not involve the shedding of mutant cfDNA.

## Acknowledgements

We thank Idit Shiff and Abed Nasseredin from the Core Research Facility at The Hebrew University Faculty of Medicine for their support in sequencing analysis, and Noa Makhervax and Dr Lilach Gavish for help in coordinating the effort. This work was supported by a generous gift from Shlomo Kramer. Supported by grants from Human Islet Research Network (HIRN UC4DK116274 and UC4DK104216 to R.S and Y.D); Ernest and Bonnie Beutler Research Program of Excellence in Genomic Medicine, The Alex U Soyka pancreatic cancer fund, The Israel Science Foundation, the Waldholtz/Pakula family, the Robert M. and Marilyn Sternberg Family Charitable Foundation, the Helmsley Charitable Trust, Grail and the DON Foundation (to Y.D), and European Research Council grant (ERC Adg 101019560) to N.F. Y.D holds the Walter and Greta Stiel Chair and Research grant in Heart studies. R.B.A received a fellowship from the Rothschild Foundation. A.R. received grant from the KAMLA Research Fund of the Hebrew University of Jerusalem and Shaare Zedek Medical Center.

## Author Contributions

Conceptualization: YD, BG, RBA, AR; Investigation: RBA, NL, EC, GF, IS, NB, DK, GK, AJ, CCS, HA, HB, NA, MMG, DV, RK, EBC, DB, TW, AQ, GC; Project administration: RBA, AR, EC; Supervision: YD, RS, BG, TK, NF; Writing – original draft: YD, RBA; Writing – review & editing: RS, BG, NF, TK, AR.

## Methods

### Clinical samples

Samples of healthy controls and hospitalized patients were obtained from patients treated in these centers: Shaare Zedek Medical Center, Jerusalem, Israel; and The Hebrew University-Hadassah Medical Center, Jerusalem, Israel. Samples of asymptomatic/mild disease were obtained from hotel-quarantined who tested positive for COVID-19. All donors serving as controls denied having any acute or major chronic illnesses or receiving any medications at the time of blood donation. Patient demographics, clinical data, and cfDNA data are detailed in **Supplementary Tables S1-S4**. Clinical parameters were extracted from patients directly or via their EMRs.

### Collection of blood samples and extraction of cfDNA

Blood samples were collected in either EDTA or STRECK tubes. For EDTA, tubes were centrifuged at 1500*g* for 10 minutes at 4°C within 4 hours of collection. Plasma was removed and recentrifuged at 3000g for 10 minutes at 4°C to remove any remaining cells. STRECK tubes were processed within 10 days of collection and were processed in the same manner, at 24°C. Plasma was then stored at −80°C until assay. cfDNA was extracted using the QIAsymphony SP instrument and its dedicated QIAsymphony Circulating DNA Kit (QIAGEN) according to the manufacturer’s instructions. DNA concentration was measured using the Qubit dsDNA HS Assay Kit (Thermo Fisher Scientific).

### Whole Genome Bisulfite Sequencing and deconvolution

Up to 75 cfDNA was subjected to bisulfite conversion using the EZ-96 DNA Methylation Kit (Zymo Research; Irvine, CA), with liquid handling on a Hamilton MicroLab STAR (Hamilton; Reno, NV). Dual indexed sequencing libraries were prepared using Accel-NGS Methyl-Seq DNA library preparation kits (Swift BioSciences; Ann Arbor, MI) and custom liquid handling scripts executed on the Hamilton MicroLab STAR. Libraries were quantified using KAPA Library Quantification Kits for Illumina Platforms (Kapa Biosystems; Wilmington, MA). Four uniquely dual indexed libraries, along with 10% PhiX v3 library (Illumina; San Diego, CA), were pooled and clustered on a Illumina NovaSeq 6000 S2 flow cell followed by 150-bp paired-end sequencing.

### Computational analysis of WGBS samples

Paired-end FASTQ files were mapped to the human genome (hg19) using bwa-meth (V 0.2.0) (32). Duplicated reads were marked by Sambamba (V 0.6.5) (33). Reads with low (<10) mapping quality, duplicated, or not mapped in a proper pair were excluded. Reads were stripped from non-CpG nucleotides and converted to PAT files using *wgbstools* (V 0.1.0) (14).

We applied our previously published fragment-level deconvolution algorithm, *UXM*(14), to estimate the cell type composition of the COVID (n=6) and control (n=6) plasma samples. The reference atlas includes 37 healthy cell types, and the features are 900 cell-type specifically unmethylated blocks. The UXM software was used with default parameters.

### Assembly of targeted DNA methylation markers

Specific CpG sites were selected by examining whole-genome bisulfite sequencing data and identifying differentially methylated or differentially unmethylated regions, having at least four CpG sites within 150 bp. We selected regions showing methylation pattern with less than 0.3 in the specific cell type and greater than 0.8 in over 90% of other cells, and designed primers to amplify ∼120 bp fragments surrounding them using the multiplex two-step PCR amplification method (34). Marker coordinates and primer sequences are provided in Supplementary table S5.

The validation of markers was done using DNA extracted from different cells and tissues, and the methylation status of the CpG block was assessed. Some markers were more sensitive if one CpG site was allowed to be methylated differently than other CpGs in the block, as indicated in Supplementary table S5.

### cfDNA PCR-sequencing analysis

cfDNA was treated with bisulfite using EZ DNA Methylation-Gold (Zymo Research) and PCR amplified with primers specific for bisulfite-treated DNA but independent of methylation status at the monitored CpG sites. Treatment with bisulfite led to degradation of 60%–90% of the DNA (on average, 75% degradation), consistent with previous reports (35). Note that while DNA degradation does reduce assay sensitivity (since fewer DNA molecules are available for PCR amplification), it does not significantly harm assay specificity since methylated and unmethylated molecules are equally affected. Primers were bar-coded using TruSeq Index Adapters (Illumina), allowing the mixing of samples from different individuals when sequencing PCR products using NextSeq sequencers (Illumina). Sequenced reads were separated by barcode, aligned to the target sequence, and analyzed using custom scripts written and implemented in R. Reads were quality filtered based on Illumina quality scores and identified by having at least 80% similarity to target sequences and containing all the expected CpGs in the sequence. CpGs were considered methylated if CG was read and were considered unmethylated if TG was read.

The fraction of unmethylated molecules in a sample was multiplied by the total concentration of cfDNA in the sample, to assess the number of specific-cell genome equivalents per milliliter of plasma. The concentration of cfDNA was measured prior to bisulfite conversion, rendering the assay robust to potential intersample fluctuations in the extent of bisulfite-induced DNA degradation.

### Plasma cfChIP-seq Chromatin analysis

Plasma collection, cfChIP assay and preprocessing of sequencing data were performed as previously described (15). Differential genes representation between hospitalized and asymptomatic samples was identified using a likelihood ratio test as described in the ‘Comparison of two groups of samples’ section of the supplementary note (15). For the enrichment test of a predefined interferon response gene set, we discarded all genes with an average TSS coverage above 10 in an independent cohort of healthy samples that was not used in this study.

### Code and data availability

Custom script for sequence analysis is available from the authors upon reasonable request. WGBS data are available at XXX.

### Statistics

To assess the significance of differences between groups, we used a 2-tailed Mann-Whitney *U* test. We calculated all CIs at the 95% level, and *P* value was considered significant when less than 0.05. In the correlation matrix, Pearson and Spearman tests were used to establish correlation between quantitative or qualitative parameters, respectively, and used FDR method to correct for multiple comparisons.

### Study approval

This study was conducted according to protocols approved by the institutional review boards at each study site: Hadassah-Hebrew University Medical Center, Jerusalem, Israel; Shaare Zedek Medical Center, Jerusalem, Israel;

Procedures were performed in accordance with the Declaration of Helsinki. Blood was obtained from donors who provided written informed consent. Case report forms were filled out by donors detailing underlying diseases.

