## Supplemental tables for "Epigenetic liquid biopsies reveal elevated vascular endothelial cell turnover and erythropoiesis in asymptomatic COVID-19 patients"

### Supplemental Figures

**Supplemental figure 1: Identification of DNA methylation markers of immune cells (A) and selected cell types from solid tissue (B).** Heatmap was derived from WGBS methylation atlas that was previously published(14) . Each row represents cell-type specific marker.

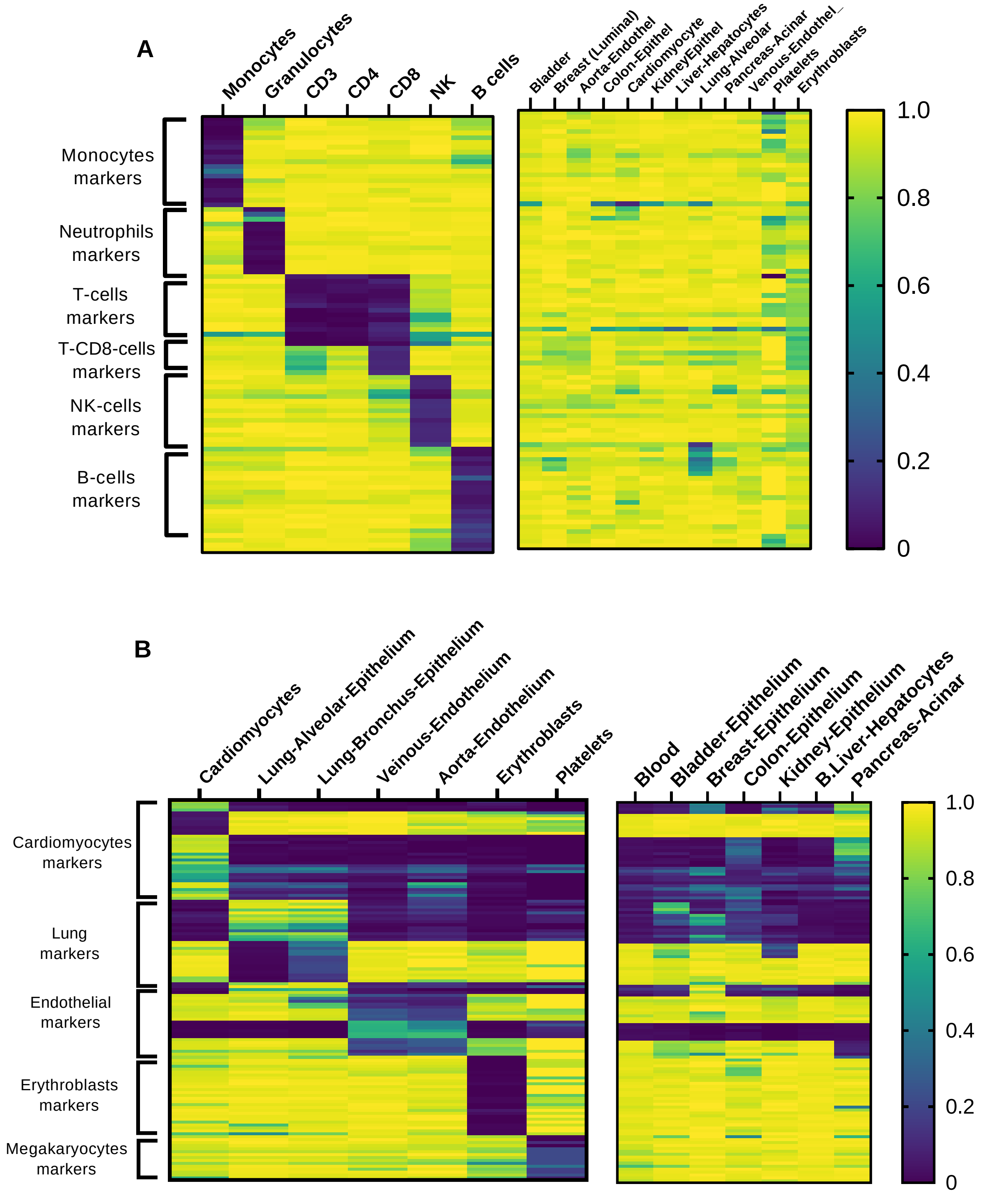

**Supplemental figure 2: Chromatin analysis of hospitalized patients compared to healthy controls.** In red, genes with differential coverage between the hospitalized and healthy samples that were used for heat map in Figure 5A.

z

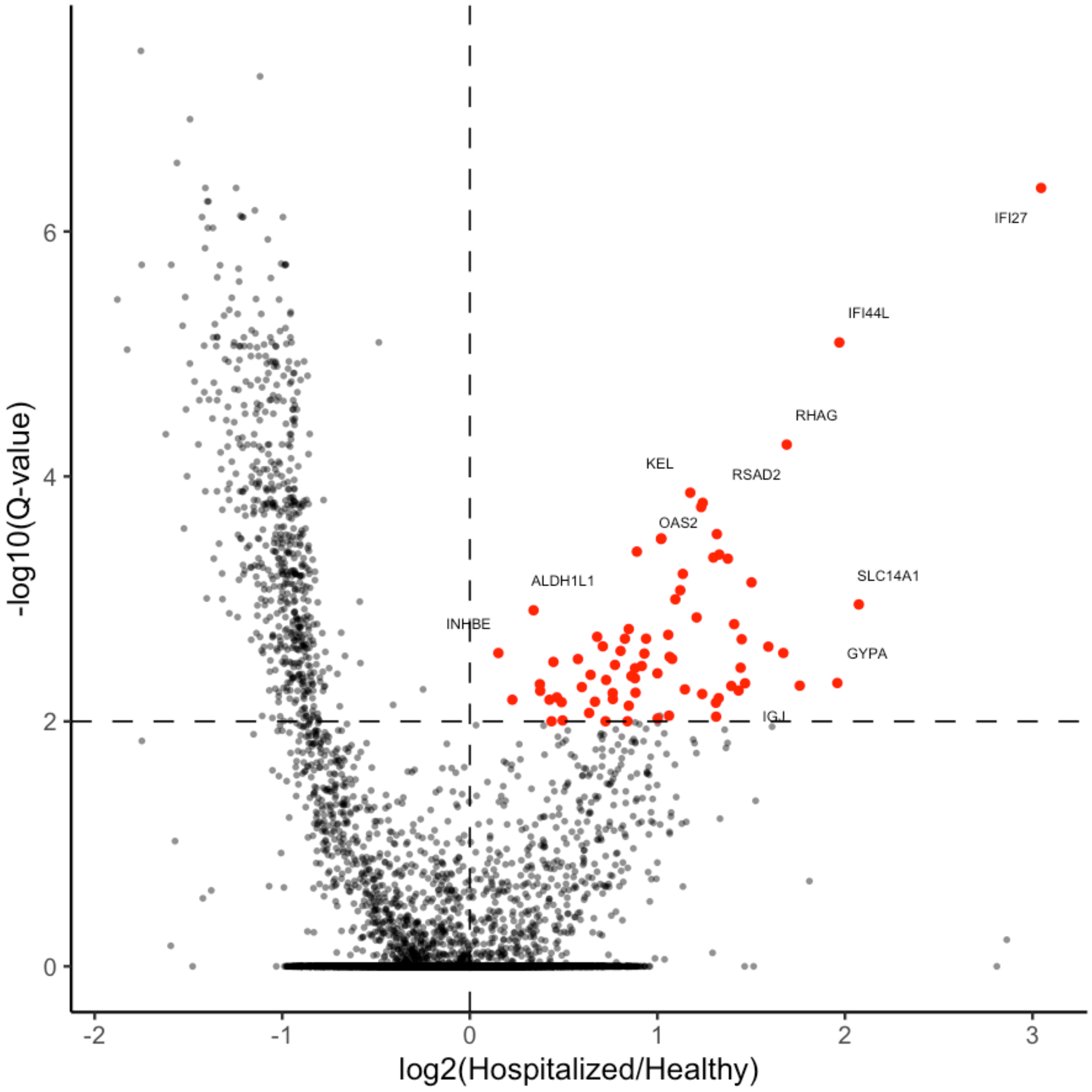

**Supplemental data**

**Supplementary table S1:** CfDNA results, demographics and characteristics of hospitalized patients.

**Supplementary table S2:** CfDNA results and demographics of asymptomatic/mild disease patients.

**Supplementary table S3:** CfDNA results and demographics of controls.

**Supplementary table S4:** CfDNA results and demographics of patients and controls (WGBS).

**Supplementary table S5:** Markers coordinates and sequences.

**Supplemental table S6:** Significantly elevated genes in hospitalized patients compared to controls.

**Supplemental table S7:** Interferon response gene set.

**Supplemental table S8:** Liver associated gene set.

**Supplemental table S9:** Erythroblasts associated gene set.

**Supplemental table S10:** Predefined interferon response gene set from the Human MSigDB Collections.
